# Functional and genetic characterization of an *incF*-type multidrug resistance plasmid isolated from fresh spinach

**DOI:** 10.1101/358887

**Authors:** Adeyinka O. Ajayi, Benjamin J. Perry, Christopher K. Yost

## Abstract

The presence of antibiotic-resistant bacteria and clinically-relevant antibiotic resistance genes within raw foods is an on-going food safety concern. It is particularly important to be aware of the microbial quality of fresh produce because foods such as leafy greens including lettuce and spinach are minimally processed and often consumed raw therefore they often lack a microbial inactivation step. This study characterizes the genetic and functional aspects of a mobile, multidrug resistance plasmid, pLGP4, isolated from fresh spinach bought from a farmers’ market. pLGP4 was isolated using a bacterial conjugation approach. The functional characteristics of the plasmid were determined using multidrug resistance profiling and plasmid stability assays. pLGP4 was resistant to six of the eight antibiotics tested and included ciprofloxacin and meropenem. The plasmid was stably maintained within host strains in the absence of an antibiotic selection. The plasmid DNA was sequenced using an Illumina MiSeq high throughput sequencing approach and assembled into contigs using SPAdes. PCR mapping and Sanger DNA sequencing of PCR amplicons was used to complete the plasmid DNA sequence. Comparative sequence analysis determined that the plasmid was similar to plasmids that have been frequently associated with multidrug resistant clinical isolates of *Klebsiella spp*. DNA sequence analysis showed pLGP4 harboured *qnrB1* and several other antibiotic resistance genes including three β-lactamases: *blaTEM-1*, *blaCTX-M-15* and *blaOXA-1*. The detection of a multidrug-resistant, clinically-relevant plasmid on fresh spinach emphasizes the importance for vegetable producers to implement evidence-based food safety approaches into their production practises to ensure the food safety of leafy greens.

## Introduction

Recent studies (6, 12, 14) have highlighted the concerns associated with the presence of clinically-relevant antibiotic resistance genes (ARGs) residing on mobile genetic elements (MGEs) isolated from foods. The spread of antibiotic-resistant bacteria and clinically-relevant ARGs from primary producers to consumers within the food chain is an emerging food safety concern (14). Consumption of fresh produce has increased due to consumer demand for healthy fresh products (15) therefore it is increasingly important to be aware of the microbial quality of minimally processed raw produce. Fresh produce such as leafy greens including lettuce and spinach are often consumed raw thereby lacking a defined kill step that would inactivate microbial pathogens (22, 23). The ability of mobile plasmids to carry and spread clinically relevant ARGs including the quinolone resistance gene family *qnr* and extended-spectrum β-lactamases within minimally processed raw foods means that a comprehensive understanding of the origin, diversity and prevalence of these plasmids is needed to help ensure food which is free from microbial threats. As part of an ongoing study on the microbial quality of fresh produce obtained from farmers’ markets, we isolated a mobile, multidrug resistance plasmid from fresh spinach, and report the genetic and functional aspects of the plasmid.

## Materials and Methods

### Spinach sampling and processing

Batches of freshly-harvested spinach were purchased on a weekly basis from a local farmers’ market between the months of June and August 2016 and transported to the laboratory under 4 °C. 150 g spinach samples were stored in individual large freezer bags containing 150 ml of buffered peptone water (BPW). The freezer bags containing BPW and spinach samples were masticated by hand for 3 min. 100 µl aliquots of the spinach peptone slurry were transferred into 5 ml test tubes containing Luria-Bertani (LB) broth amended with 0.8 µg ciprofloxacin/ml. Ciprofloxacin was used to enrich for ciprofloxacin-resistant bacteria that may harbour fluoroquinolone resistance plasmids. Enrichment cultures were incubated overnight in a shaking incubator at 37 °C at 200 x *g*.

### Plasmid isolation using a conjugation approach

Antibiotic resistance plasmids were isolated using a bacterial conjugation approach. Overnight spinach enrichment cultures were used as donor cells with a sodium azide-resistant *Escherichia coli* strain J53 recipient strain. 1 ml of donor and recipient cultures were centrifuged at 12,000 rpm for 3 min, washed twice with LB broth to remove ciprofloxacin or sodium azide from the mating bacterial suspension and resuspended in 100 µl of LB broth. 100 µl of donor cells were mixed with 100 µl of recipient cells and the mixture including 100 µl of donor and recipient controls were spot-plated on MacConkey agar plates and incubated overnight at 37 °C. MacConkey agar was used to select for a donor population of relevant enteric bacteria. Bacterial matings were scrapped off and resuspended in 900 µl of sterile water. 100 µl aliquots of the resuspension were plated onto MacConkey agar plates amended with 150 µg sodium azide/ml and 0.8 µg ciprofloxacin/ml to select for transconjugants. Transconjugant plates were incubated overnight at 37 °C and resulting *E. coli* J53 transconjugant colonies were screened for the presence of antibiotic resistance plasmids using Eckhardt gel electrophoresis.

### Multidrug resistance experiments and restriction digests

*E. coli* J53 transconjugant colonies were screened for multidrug resistance using Luria-Bertani (LB) agar media supplemented individually with antibiotics at the following concentrations: chloramphenicol (30 µg/ml), erythromycin (200 µg/ml), tetracycline (10 µg/ml), spectinomycin (100 µg/ml), ampicillin (100 µg/ml), kanamycin (50 µg/ml), ciprofloxacin (0.8 µg/ml) and meropenem (4 µg/ml). An *E. coli* J53 transconjugant that was resistant to multiple antibiotics (ciprofloxacin, tetracycline, ampicillin, kanamycin, meropenem and spectinomycin) was selected for further study and plasmid DNA was isolated using a Qiagen Miniprep kit as per the manufacturer’s protocol (Hilden, Germany).

DNA restriction digest with multiple restriction enzymes was performed to estimate the size of the multidrug resistance (MDR) plasmid.

### Conjugation assay using *Pantoea agglomerans* 5565

Conjugation assays were also carried out using the *E. coli* J53 transconjugant as the donor strain and a rifampicin-resistant *Pantoea agglomerans* 5565 as the recipient strain. For conjugation assays using *E. coli* J53 transconjugants and *P. agglomerans* 5565, 1 ml of donor and recipient cultures from overnight cultures were centrifuged at 12,000 x *g* for 3 min, washed twice with LB broth and resuspended in 100 µl of LB broth. 100 µl of donor cells were mixed with 100 µl of recipient cells and the mixture including 100 µl of donor and recipient controls were spot-plated on LB agar plates and incubated overnight at 37 °C. Bacterial matings were scrapped off the plates and resuspended in 900 µl of sterile water. 100 µl aliquots of the resuspension were plated onto Vincent’s minimal media (VMM) agar plates (26) amended with 0.8 µg ciprofloxacin/ml and 100 µg rifampicin/ml to select for *P. agglomerans* transconjugants as *P. agglomerans* 5565 can grow in minimal media conditions. Transconjugant plates were incubated overnight at 37 °C and resulting transconjugant cells were screened for the presence of the multidrug resistance plasmid by Eckhardt gel electrophoresis. Conjugation transfer efficiency was calculated as the number of transconjugants per donor cell.

### Heavy metal resistance assay

The functionality of the predicted heavy metal resistance gene region encoded on pLGP4 was tested using the broth micro-dilution method of minimum inhibitory concentration (MIC) as described by EUCAST (9). We modified this procedure for heavy metal resistance testing. The modified procedure is briefly described; pLGP4-bearing *E. coli* J53 transconjugant and an *E. coli* J53 without the plasmid were sub-cultured in LB broth and grown overnight, shaking at 200 x *g* for 16 h at 37 °C. Copper and silver were selected as representative metals and the highest tested concentrations were 3mM and 2.5mM for copper and silver respectively. A 96-well plate was used in the MIC determination. 100 µl of LB broth was added to columns 2 to 12. LB broth was not added to well 1. 200 µl of copper (CuSO_4_) and silver (AgNO_3_) stock solutions 6mM and 5mM respectively were added to well 1. Subsequently, in a two-fold serial dilution, 100 µl of copper and silver stocks from well 1 were transferred to well 2 up till well 11, in each instance, the samples were mixed thrice by pipetting aseptically. A metal-free control was included in column 12, hence 100 µl was aspirated out of well 11. From overnight cultures of the pLGP4-bearing *E. coli* J53 transconjugant and the non plasmid-carrying *E. coli* J53, 100 µl of 10^6^ cfu/ml concentrations of the cultures were added to the 12 wells in each row to give a 10^5^ cfu/well. The experiment was conducted in triplicate. The plates were incubated in a BioTek Gen5 plate reader machine (version 3; BioSPX, Belgium) for 24 h at 37 °C and OD_600_ values were read at every 4 h interval.

### Plasmid stability assay

To determine the maintenance of the MDR plasmid in bacterial cells in the absence of antibiotic selection, plasmid stability was tested in Luria-Bertani (LB) broth. Single colonies of both *E. coli* J53 and *P. agglomerans* pLGP4 transconjugants were individually inoculated into 5 ml LB broth without antibiotic and grown overnight at 37 °C with agitation. Serial dilutions were plated in duplicate on LB agar and LB agar with ciprofloxacin (0.8 µg/ml) to enumerate total viable cells and plasmid-containing cells respectively. 100 µL of the undiluted bacterial cells were transferred into fresh 5 ml LB broth without antibiotic on consecutive days for up to 30 days, enumerations of total viable cells and plasmid-containing cells were performed on days 2, 4, 8, 16, 24 and 30. This approximated 2,160 bacterial generations, based on a generation time of 20 min.

### High throughput DNA sequencing of the plasmid

The plasmid DNA was isolated from the *E. coli* J53 pLGP4 transconjugant and purified using a Qiagen plasmid Miniprep kit (Hilden, Germany). 2 ng of the plasmid DNA was used as a starting material for library preparation using the Illumina Nextera XT DNA Library Preparation Kit according to the manufacturer’s protocols (San Diego, 92122, CA, USA). The prepared pLGP4 DNA library was multiplexed into a single 300 bp paired-end sequencing run on an Illumina MiSeq using a 600 cycle MiSeq v3 Reagent Kit. The resulting paired-end sequences were used for assembly using SPAdes (4) and assembled contigs were visualised using Bandage (25). Contig Assembly using Rearrangements (CAR) genome reference-assisted scaffolding tool (16) was used to order the plasmid contigs into a single scaffold. PCR mapping with primers designed using Snapgene (GSL Biotech LLC, Chicago, USA) and Sanger DNA sequencing of PCR amplicons were used to close the gaps between contigs and complete the full DNA sequence of the plasmid. The completed plasmid sequence was imported into the Rapid Annotation for Subsystem Technology (RAST) server (3) for gene prediction and annotation. The comparative analysis of pLGP4 with other closely-related plasmids of clinical and environmental origins was done using the CGview comparative tool hosted on the CGview server (10) and Mauve (8). Phylogenetic alignments and the construction of gene trees were done using Geneious (version 9.1.0; Biomatters, U.S.A). Gene synteny diagrams were drawn using Easyfig (http://mjsull.github.io/Easyfig/). The plasmid sequence was imported into the Plasmid Replicon Finder server (7) for replicon typing. The sequence of the assembled plasmid (pLGP4) was deposited into the NCBI database under the accession number MF116002.1 (https://www.ncbi.nlm.nih.gov/nuccore/MF116002).

## Results and Discussion

pLGP4 is a 198, 388 bp, 51.7% G-C, multidrug-resistant *incF* plasmid that has three replicon regions (*repFII*, *repFIA* and *repFIB*) which are conserved in several *incF* plasmids. A study reported that the multi-replicon property of most *incF* plasmids ensures their stable inheritance in a broader range of bacterial hosts and helps to avoid incompatibility issues with other plasmids of the same incompatibility group (19, 24). Multi-replicon *incF* plasmids can use either *repFII*, *repFIA* or *repFIB* genes for replication initiation hence these multi-replicon *incF* plasmids can switch between *rep* genes which enables multiple *incF* plasmids to reside within the same cell (19, 24).

pLGP4 showed resistance to six of the eight tested antibiotics and included: kanamycin, tetracycline, ampicillin, spectinomycin, ciprofloxacin and meropenem. The plasmid did not confer resistance to chloramphenicol and erythromycin. pLGP4 harboured the quinolone resistance gene *qnrB1* gene which has been associated with clinical isolates of pathogenic *Klebsiella* species (13). *K. pneumoniae* belongs to the group of pathogens often referred to as the ES-K-A-P-E pathogens based on their clinical importance and antibiotic resistance (others are *Escherichia coli*, *Acinetobacter baumannii*, *Pseudomonas aeruginosa* and *Enterococcus faecalis*) (5) The plasmid also encoded three different classes of β-lactamase genes: *blaOXA- 1*, *blaTEM-1* and the carbapenem-hydrolysing extended spectrum β-lactamase gene (*blaCTX-M-15*) that were closely related to respective β-lactamases found on plasmids from antibiotic-resistant clinical pathogenic isolates (Fig. 2B). *IncF* plasmids are a clinically relevant group of plasmids commonly implicated in the dissemination of *blaCTX-M-15* genes (2, 13, 19). Carbapenems are regarded as the “last line” antibiotics due to their effectiveness in the treatment of many multiple antibiotic-resistant infections (27). Fluoroquinolones and β-lactam antibiotics are two of the most powerful class of antibiotics used in clinical settings. These antibiotics have broad spectrum applications in the treatment of a wide range of Gram-positive and Gram-negative bacterial infections (21, 27), hence resistance to these antibiotics are a public health concern. pLGP4 harboured the tetracycline efflux gene (*tetA*) and its transcriptional regulator (*tetR*), the fluoroquinolone-modifying, aminoglycoside N(6’) acetyltransferase gene (*aac(6’)-lb*), an aminoglycoside N(3’) acetyltransferase, two aminoglycoside phosphotransferase gene variants (*strA* and *strB*) known to confer resistance to streptomycin, a trimethoprim-sulfamethoxazole resistance gene (*dfrA1*) flanked by an integron-integrase gene (*intl1*) that encodes the clinically-important class 1 integrons and are usually associated with the dispersal of *dfrA1* genes. The integron-integrase gene encoded on pLGP4 has been determined to be a novel variant and has since been curated into the INTEGRALL integron database, with the designation In1423 (http://integrall.bio.ua.pt/?acc=MF116002) (17). pLGP4 also harboured the dihydropteroate synthase gene (*sul1*) known to confer sulphonamide resistance (Fig. 1A). A transposon-insertion within a chloramphenicol acetyltransferase gene (*catB*) provides an explanation for the lack of chloramphenicol resistance.

**Figure 1A.**
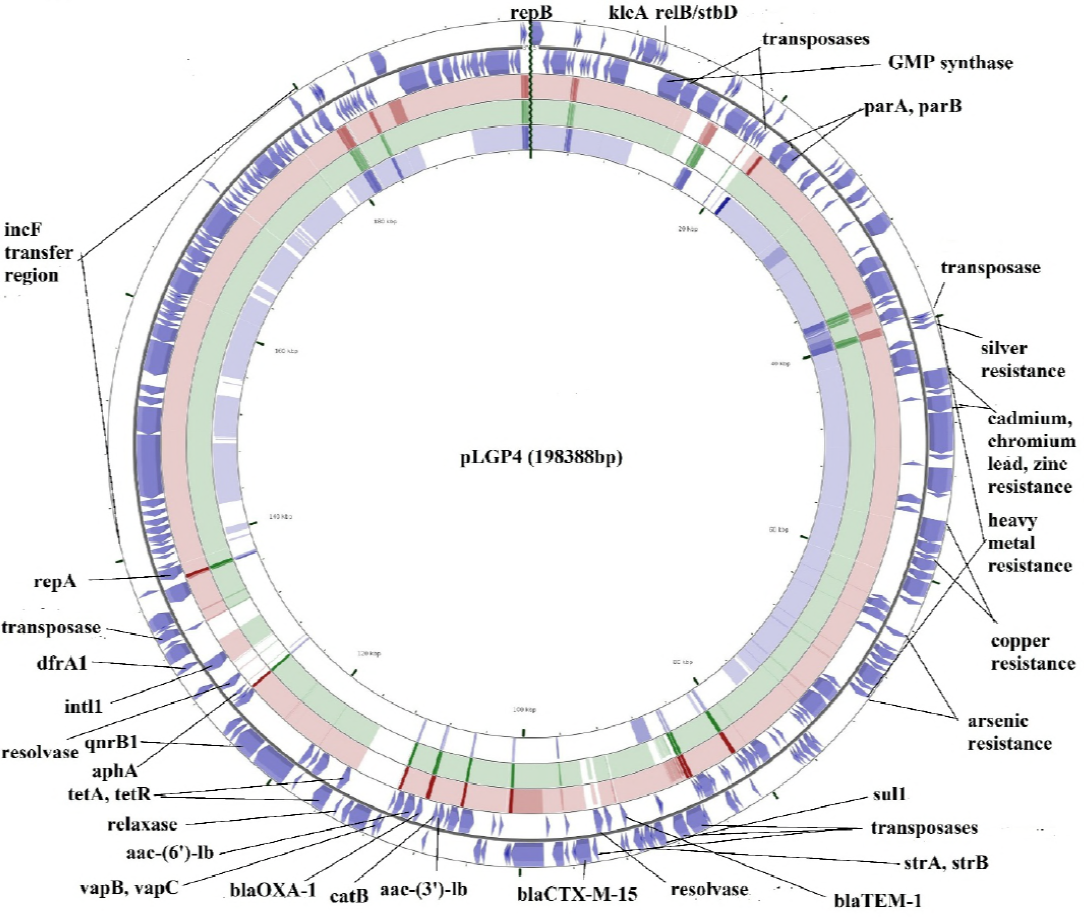
Illustrative map comparing pLGP4 (outer two rings with blue arrows) with three closely-related plasmids: p6234, pKPSH and pKPN3. The pink, green and blue regions correspond to p6234 (body fluids, USA), pKPSH11 (WWTP biosolids, Israel) and pKPN3 (blood sample, Colombia) respectively. The heights of each arc in the corresponding BLAST-hit regions are proportional to the percent identity of the hits. Overlapping hits appear as darker arcs.

The plasmid encodes a large metal resistance module (33 kbp) (Fig. 1A) that contained a cluster of six genes putatively encoding for arsenic resistance, a silver-binding gene and 10 genes putatively encoding copper, lead, cadmium, zinc and mercury resistance. In the presence of copper at concentrations of 1.5mM and 3mM, the pLGP4 –carrying *E. coli* J53 had a consistent and uniform increase in growth with high peak OD_600_ values (Fig. 3). *E. coli* J53 without the plasmid grew steadily at 1.5mM copper, but its growth was significantly hindered at 3mM copper (ANOVA; p < 0.05) (Fig. 3). In the presence of silver at a concentration of 1.25mM, the pLGP4 –carrying *E. coli* J53 had a steady and uniform growth increase (Fig. 4). The pLGP4-carrying *E. coli* J53 began to grow steadily at 2.5mM silver after 12 hrs of incubation (Fig. 4). *E. coli* J53 without the plasmid grew steadily at 1.25mM silver, but its growth was significantly hindered at 2.5mM silver (ANOVA; p < 0.05) (Fig. 4). The results of the copper and silver resistance assays suggest the copper and silver resistance genes in pLGP4 are functional.

**Figure 1B.**
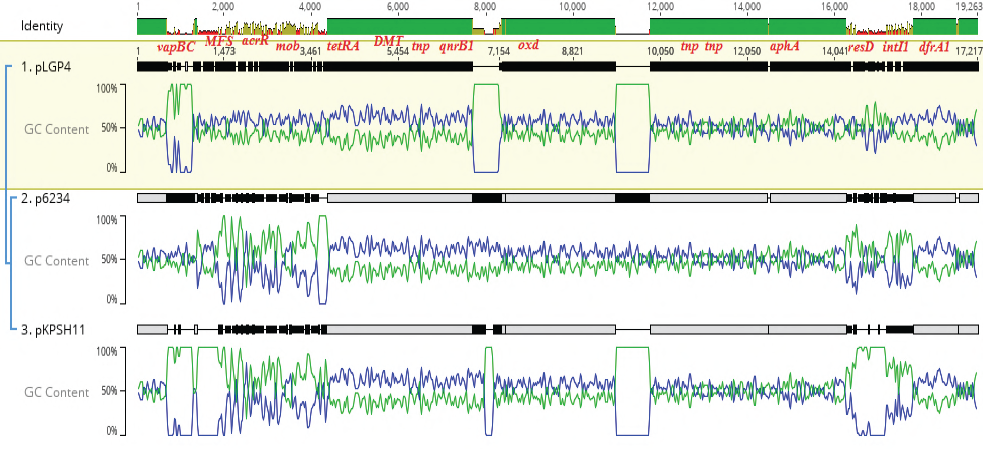
Comparative sequence alignment analysis showing the portion of pLGP4 sequence with marked genetic variation to corresponding portions of the other closely- related pKPN3-like plasmids: p6234 and pKPSH11. Green alignment bars indicate 100% identity with BLAST hits, greenish-brown bars signify 30-100% identity with BLAST hits while red bars indicate an identity less than 30% to BLAST hits. The blue waves show the GC content while the green waves represent the AT content. The *vapBC* toxin-antitoxin genes, the membrane efflux transport gene *MFS* and the Tn3-type resolvase *resD* are all unique to pLGP4. The *tet* family transcriptional repressor gene *acrR*, the relaxase gene *mob* and the integron integrase gene *intI1* all show some variation in their genetic sequence when compared to those found in p6234 and pKPSH11.

**Figure 1C.**
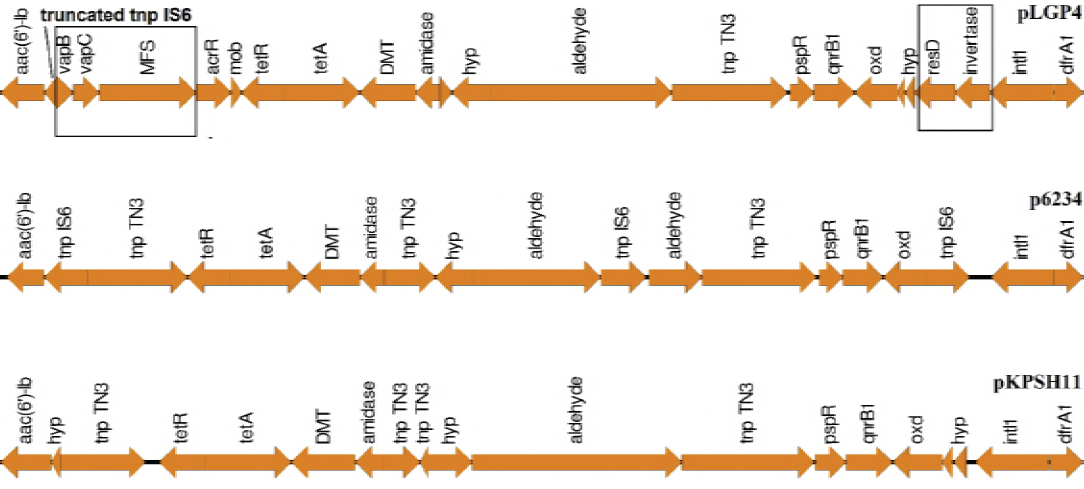
Gene synteny analysis highlighting the portion of the pLGP4 sequence showing some evidence of genetic variance when compared to the other pKPN3-like plasmids. The addiction genes *vapBC*, a flanking membrane transport *MFS* gene (could be reason for *vapBC* mobilization), a resolvase gene *resD* and a DNA invertase gene are all unique to pLGP4. These unique and varied features of pLGP4 show some evidence of variation due to evolutionary divergence between the plasmids. The mobilization of the *qnrB1* gene in the three plasmids may have been due to the action of Tn3-type transposases (*tnp* TN3).

**Figure 1D.**
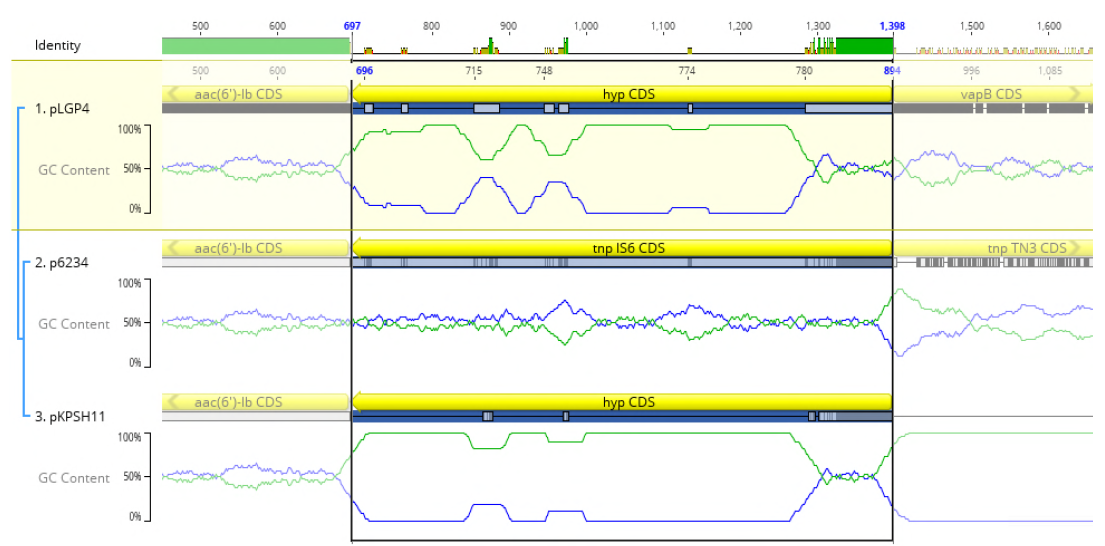
Comparative sequence alignment analysis of the flanking region of the vapBC genes. The *hyp* gene (hypothetical protein) flanking the vapBC genes shows some similarity to an IS6 family transposase (*tnp* IS6) of pKPN3, suggesting this *hyp* gene may be a remnant of the *tnp* IS6 gene. Green alignment bars indicate 100% identity with BLAST hits, greenish brown bars signify 30-100% identity with BLAST hits while red bars indicate an identity less than 30% to BLAST hits. The blue waves show the GC content while the green waves represent the AT content.

**Figure 2A.**
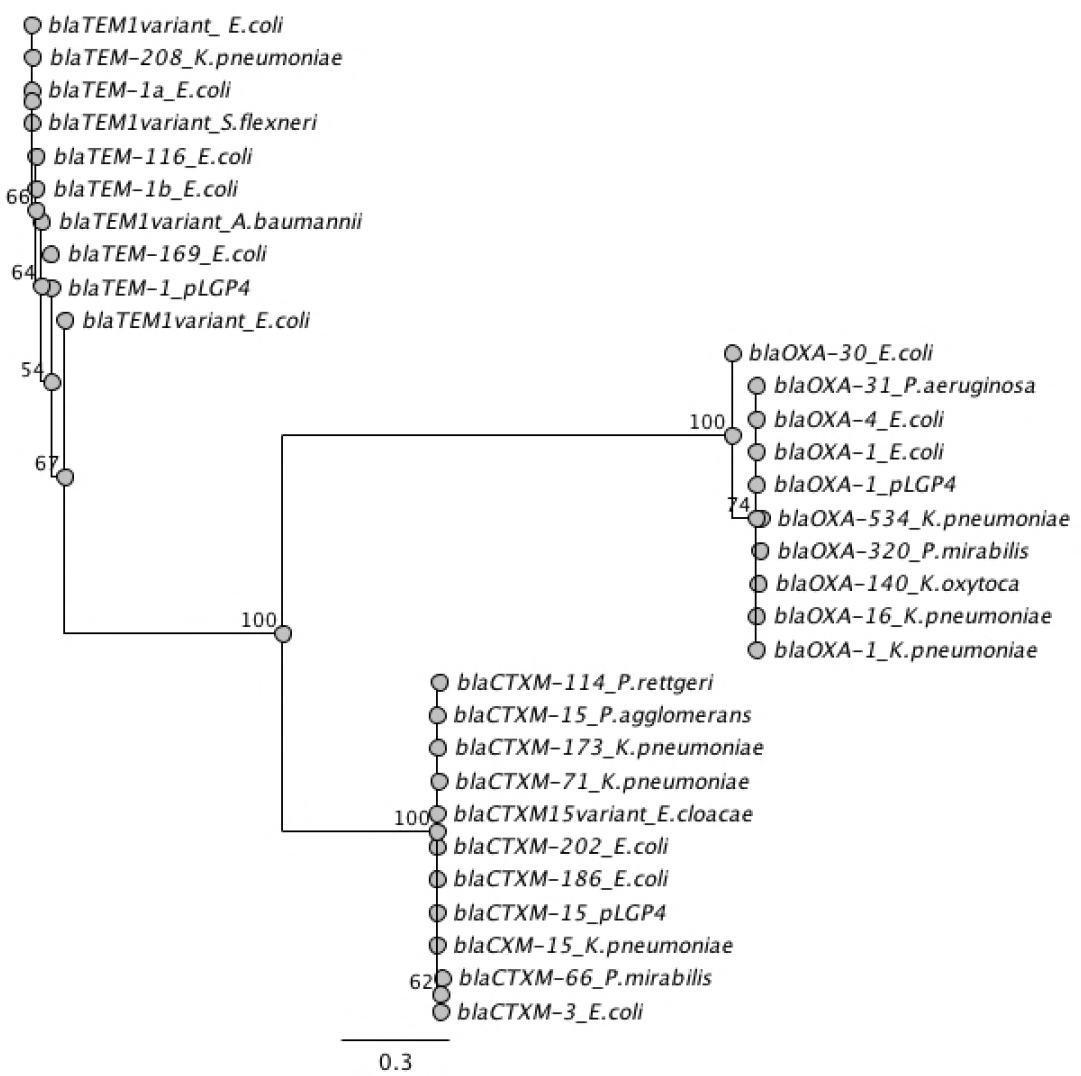
Gene phylogeny of the pLGP4-encoded β-lactamase genes (*blaOXA-1*, *blaTEM-1* and *blaCTXM-15*) with respective β-lactamase genes found in clinical pathogenic isolates. The β-lactamase genes found in K. pneumoniae were all encoded on mobile plasmids.

**Figure 2B.**
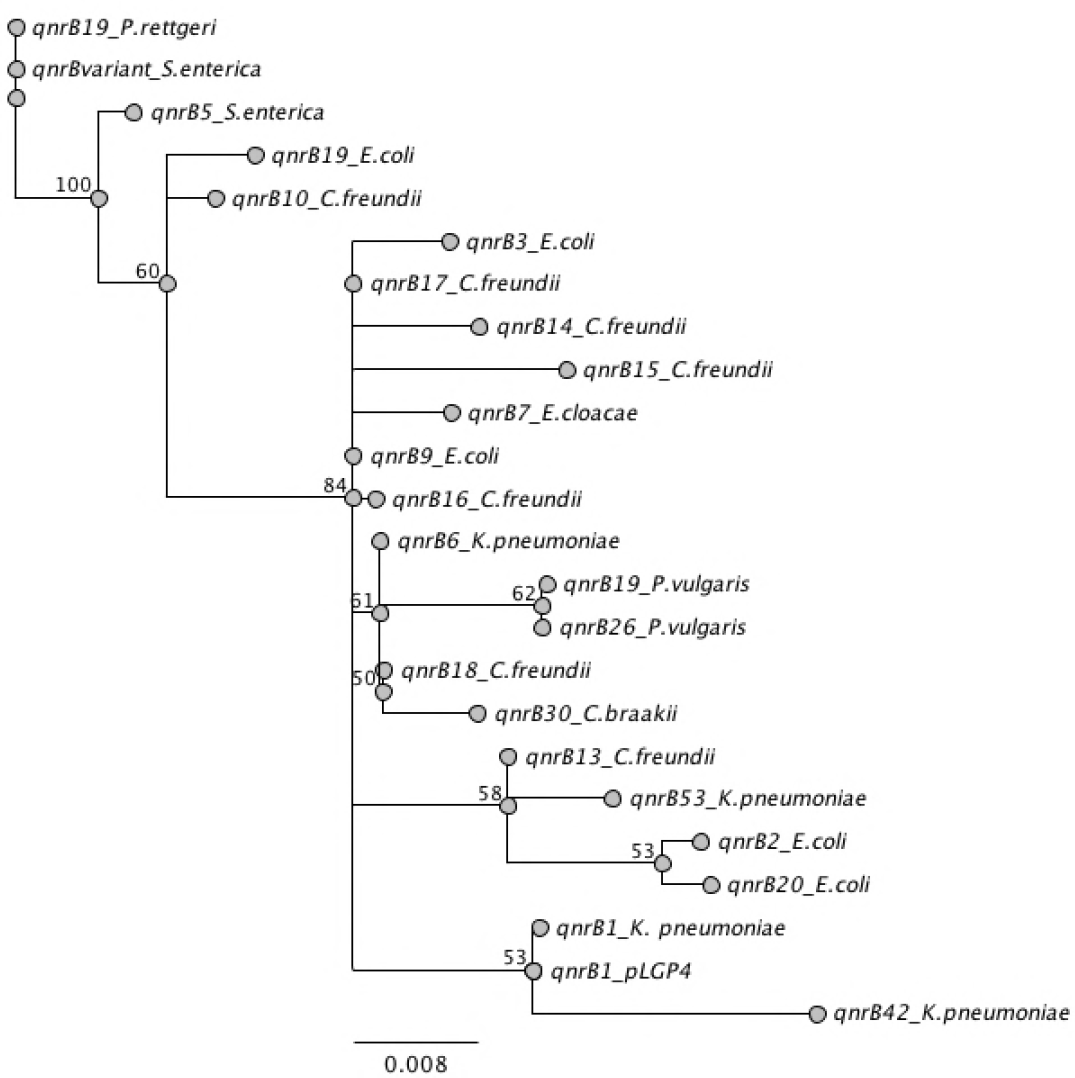
Gene phylogeny of the pLGP4-encoded *qnrB1* gene with *qnrB* variants found in clinical pathogenic isolates. The *qnrB* variants found on *K. pneumoniae* were all encoded on mobile plasmids. The pLGP4-encoded *qnrB1* was most closely-related to a qnrB1 gene found in a plasmid pMRKp1 isolated from a clinical isolate (urinary tract infection) of *K. pneumoniae.*

**Figure 3:**
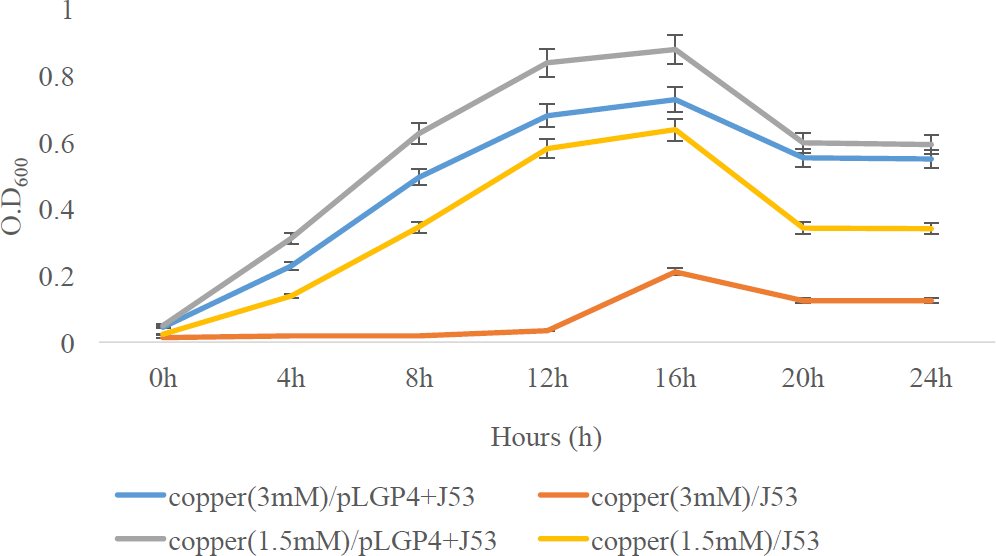
OD_600_ absorbance trends of copper/pLGP4+J53 and copper/J53 combinations over a 24 h period. 3mM copper (red line) significantly hindered the growth of *E. coli* J53 without plasmid pLGP4 (p < 0.05). The pLGP4-carrying *E. coli* J53 had a consistent growth increase in the presence of 3mM copper (blue line).

**Figure 4:**
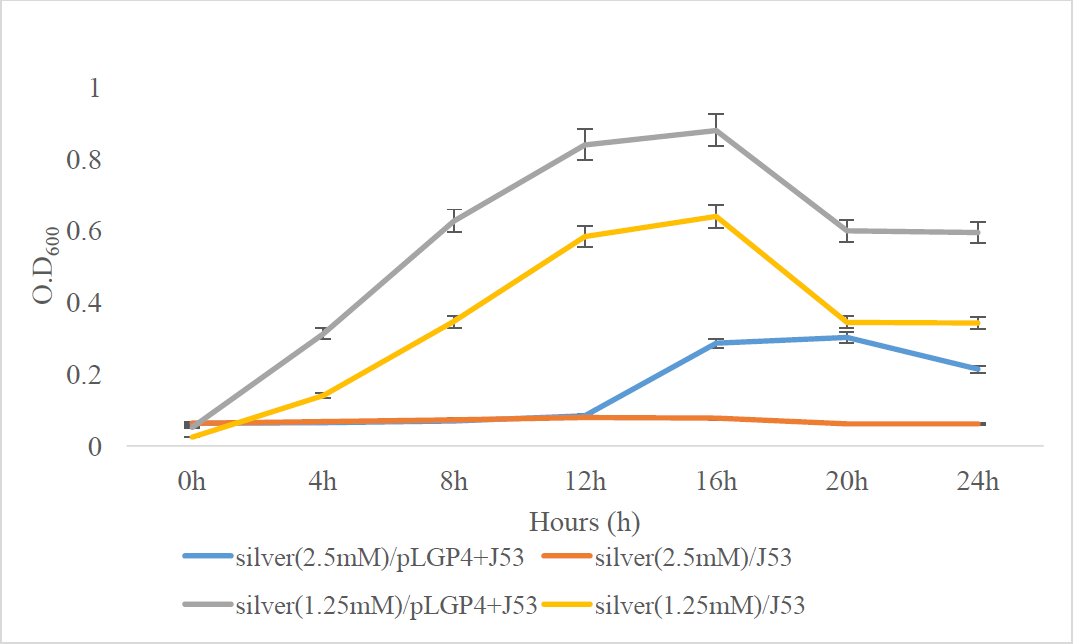
OD_600_ absorbance trends of silver/pLGP4+J53 and silver/J53 combinations over a 24 h period. 2.5mM silver (red line) significantly hindered the growth of *E. coli* J53 without plasmid pLGP4 (p < 0.05). The pLGP4-carrying *E. coli* J53 began to have a steady growth increase in the presence of 2.5mM silver (blue line) after 12 h of incubation.

**Figure 5.**
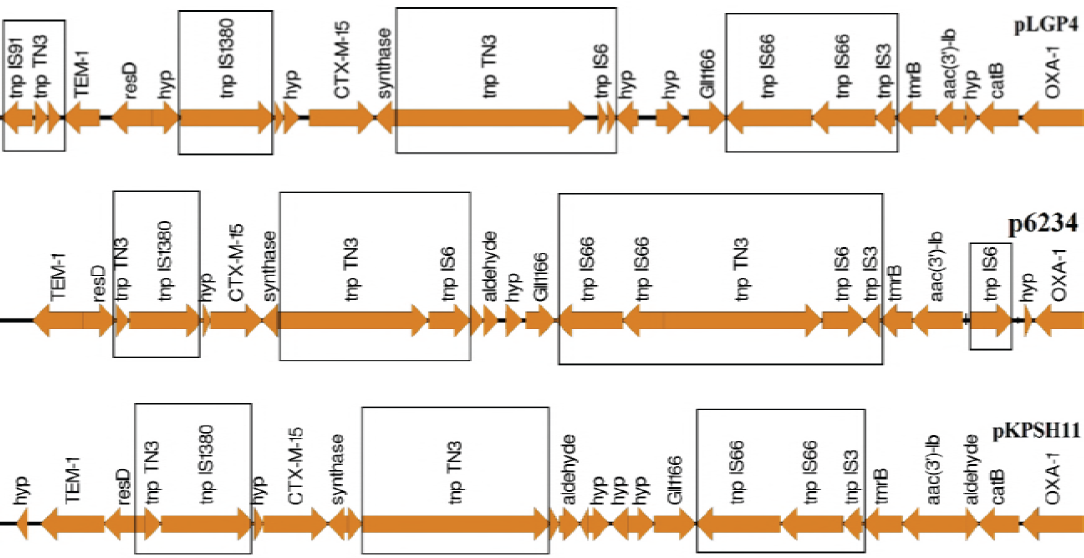
Gene synteny analysis of the flanking regions of the three β-lactamases (TEM1, CTX-M-15 and OXA-1). This gene synteny analysis shows the presence of Tn3-type transposases (*tnp* TN3) and other IS-family transposases (IS1380, IS66, IS6, IS3, IS91) within the flanking regions of the three β-lactamases (TEM1, CTX-M-15 and OXA-1) encoded in the three closely-related pKPN3-like plasmids. Other genes within the flanking regions include: *resD* (resolvase), *hyp* (hypothetical proteins), *aac (3′)-lb* (aminoglycoside acetyltransferase), *catB* (chloramphenicol acetyltransferase) and *tmrB* (tunicamycin resistance gene).

pLGP4 also carried several genes known to encode mobile resistance elements including Tn3- type and IS family transposases which are important in the mobilization of clinically-relevant antibiotic resistance genes (18), and resolvases (important in multimer resolution and plasmid stability) (Fig. 5). The quinolone and β-lactam resistance genes of pLGP4 were similar to those found in antibiotic-resistant clinical pathogenic isolates (Figs. 2A–2B). The close similarity between the fluoroquinolone and β-lactamase genes encoded on pLGP4 and those found in antibiotic-resistant clinical pathogenic isolates suggests these antibiotic resistance genes may be able to confer antibiotic resistance at clinically relevant levels should a pathogen acquire pLGP4 via conjugation.

pLGP4 was shown to be a functional conjugative plasmid and transferred to *P. agglomerans* 5565 at a transfer frequency of 3.7 x 10^−4^ per donor cell. pLGP4 encoded a cluster (40 kb) of 25 *incF* conjugative transfer (*tra* and *trb*) genes that are conserved among several *incF* plasmids. The results of the stability assays showed persistence of the plasmid in the two host strains in the absence of antibiotic selection. 93.8% (9.38 log_10_ at day 1 to 8.8 log_10_ at day 30) of the total population maintained the pLGP4 plasmid over a 30-day test period (2,160 bacterial generations) in *E. coli* J53 while 97.1% (9.48 log_10_ at day 1 to 9.2 log_10_ at day 30) of the total population maintained pLGP4 over a 30-day test period (2,160 bacterial generations). There was a slight reduction in the numbers of plasmid containing *E. coli* J53 and *P. agglomerans* cells, this reduction was not significant (ANOVA; p < 0.05). pLGP4 encoded an anti-restriction methylase gene (*klcA*) and two different types of addiction modules: *relB/stbD*, *vapC/vapB*) (Fig. 1A). pLGP4 also codes for plasmid partitioning proteins *(parA* and *parB*) (Fig. 1A) important for plasmid stability as persistence of pLGP4 in *E. coli* J53 and *P. agglomerans* without selective pressure suggests the pLGP4- encoded partitioning and addiction genes are functional.

pLGP4 has a 99% identity over 92% query coverage with pKPSH11 and a 100% identity over a 92% query coverage with p6234. It has a 99% identity over a 59% query coverage with pKPN3 according to BLASTN analysis (1). The conserved nature of the backbone region of pLGP4 among the *Klebsiella pneumoniae*-associated pKPN3-like plasmid family suggests this plasmid is related to a common plasmid ancestor along with other members of the pKPN3-like plasmid family (Fig. 1A, Fig. S1). pKPN3 is distantly-related to pKPSH, p6234 and pLGP4 (Fig. 1A, Fig. S1). These pKPN3-like plasmids were characterized by a distinct structural arrangement, having an accessory module that contained all the antibiotic resistance genes and the backbone module that contained the plasmid conjugative transfer, replication and maintenance genes. While the backbone module is highly conserved among members of the pKPN3-like plasmid family (Fig. 1A), the accessory modules of the plasmids are similar but not identical.

The original bacterial host of pLGP4 is currently unknown. However, this plasmid belongs to the family of pKPN3-like plasmids that are frequently associated with different strains of *K. pneumoniae* (13). BLASTN analysis showed that the *K. pneumoniae*-associated pKPN3-like plasmids were closely-related to each other despite their geographical separation- pKPN3 (Colombia), pKPSH11 (Israel) and p6234 (USA). Comparative analysis revealed some evidence of genetic variation between the plasmids (Figs. 1B–1C, Fig. S1). These variations may have arisen from different selection pressures that enabled the plasticity (acquisition and loss) of these genetic elements. pLGP4 encoded unique genes: toxin-antitoxin genes *vapB* and *vapC*, inner membrane efflux gene *MFS*, Tn3 resolvase *resD* and a DNA invertase gene (Fig. 1C). *vapBC* was flanked by a gene encoding for a hypothetical protein that was similar to a truncated *IS*6 family transposase (Figs. 1C–1D). The truncated IS6 family transposase gene *tnp* IS6 may have been responsible for the mobilization of *vapBC* and this mobilization event may have led to the disruption of the transposase (*tnp IS6*) gene (Figs. 1B–1D).

The ability of mobile, multidrug resistance plasmids to be easily mobilized to other bacterial species means the origin of pLGP4 cannot be accurately ascertained using comparative genomics. The properties of this plasmid suggest it may constitute a threat to food safety and public health if carried by bacteria capable of colonizing the human gut or transferred to pathogens residing on produce that is intended to be consumed raw.

